# UdonPred: Untangling Protein Intrinsic Disorder Prediction

**DOI:** 10.64898/2026.01.26.701679

**Authors:** Julius Schlensok, David Wagemann, Tobias Senoner, Markus Haak, Burkhard Rost

## Abstract

**Motivation:** Regions in intrinsic disordered proteins (IDPs) constitute important continuous aspects of protein function. While their existence on a structural continuum is widely accepted, most computational predictions have, nevertheless, focused on binary classifications. Existing datasets are severely limited in size and experimental evidence for continuous disorder.

**Results:** Building on recently released datasets of continuous protein disorder and flexibility, we introduce UdonPred, a lightweight neural network exclusively inputting embeddings from the protein Language Model (pLM) ProstT5 to predict per-residue protein disorder from sequence alone. Training and evaluating UdonPred on seven datasets with divergent definitions of disorder and flexibility suggests that not model capacity, but agreement and nuance of disorder annotations, remains the main driver of performance. Binary disorder annotations can be reliably predicted from a multitude of different disorder and flexibility datasets, but there is still room for improvement in predicting continuous disorder.

**Availability:** All code and data used for training and evaluation is available under an open-source license at https://github.com/davidwagemann/udonpred.

**Supplementary information:** Supplementary data are available at *Journal Name* online.

## 1. Introduction

Intrinsically disordered proteins (IDPs) and regions therein (IDRs), which are highly abundant in all domains of life and especially in eukaryotes (Pancsa and Tompa, 2012; Peng et al., 2015; DeForte and Uversky, 2016), defy the classical structurefunction paradigm by performing their biological roles without adopting a single, stable three-dimensional structure under physiological conditions (Wright and Dyson, 1999; Dunker et al., 2001). This conformational plasticity allows IDPs to participate in crucial cellular processes that rely on dynamic interactions, such as signal transduction, transcriptional regulation, and cell-cycle control (Dunker et al., 2008; Babu, 2016; Uversky, 2021; Bondos et al., 2022). IDPs are strongly associated with disease in humans (Uversky, 2009, 2014; Babu et al., 2011) and their misfunction and aggregation are hallmarks of numerous conditions, including cancer (Iakoucheva et al., 2002), cardiovascular diseases (Cheng et al., 2006), neurodegenerative disorders (Uversky, 2009; Uversky et al., 2014), and mental disorders (Zhang et al., 2024); they are an important target in drug development (Ruan et al., 2019; Lazar et al., 2025). With experimental annotations still sparse (Nugnes et al., 2025; Piovesan et al., 2022b), the accurate prediction of protein disorder remains a fundamental task in modern bioinformatics.

Disorder predictions use ever more sophisticated machine learning techniques and are trained on increasingly diverse feature sets (Piovesan et al., 2022c; Pang and Liu, 2024; Wang et al., 2024; Kotowski et al., 2025; Liu et al., 2025). Since the success of SETH (Ilzhöfer et al., 2022), most state-of-the-art disorder prediction methods incorporate embeddings from protein Language Models (pLMs) (Mehdiabadi et al., 2026), such as ESM-2 (Lin et al., 2023) and ProtT5 (Elnaggar et al., 2022) - large pre-trained models that learn to extract important nonlocal information about proteins from large unlabeled sequence databases (Ilzhöfer et al., 2022; Mehdiabadi et al., 2026; Kabir and Hoque, 2024). Manually verified disorder predictions have just been allowed as evidence in the newest release DisProt database, highlighting the progress of the whole field Nugnes et al. (2025).

High-fidelity experimental data of protein disorder remains very limited. For instance, only about 3% of all the proteins in the Protein Data Bank (PDB; Berman et al., 2000) contain IDRs by any definition (Aspromonte et al., 2024; Xue et al., 2012), and fewer yet contain more stringent IDRs by the definition that has at least every third human protein have an IDR (Galea et al., 2009). Experimental annotations for intrinsic disorder are derived using a variety of techniques, including missing electron density in X-ray crystallography (Dunker et al., 2001), Nuclear Magnetic Resonance (NMR) spectroscopy (Sormanni et al., 2017), Circular Dichroism (CD; Dunker et al., 2001), and small-angle X-ray scattering (SAXS; Gräwert and Svergun, 2020). Each of these captures a different aspect of protein disorder across different timescales (Sormanni et al., 2017). Disorder annotations are spread across multiple databases, each curating a different, often nonoverlapping sample of the IDP/IDR population based on specific structural or functional criteria and with inherent biases from curation and experimental techniques (Necci et al., 2018; Piovesan et al., 2022b, 2023).

This fragmentation is compounded by inconsistencies in how disorder is defined and represented. The majority of available annotations used to train most models are binary, classifying residues simplistically as “ordered” or “disordered” (Nielsen and Mulder, 2019; Aspromonte et al., 2024; Hsu et al., 2020). This paradigm fails to capture the more nuanced continuum of protein disorder, where disordered regions can exist in disordered and ordered states depending on experimental conditions, bind in fuzzy complexes, or undergo a disorder-to-order transition upon binding (Hsu et al., 2020; Piovesan et al., 2022a). The CheZOD scoring scheme (Nielsen and Mulder, 2020), which derives continuous disorder scores from NMR chemical shifts, was the first resource of continuous disorder annotations, but was never widely adopted, partially due to being rather small (Nielsen and Mulder, 2019). Prediction methods have only recently been trained on continuous annotations (Ilzhöfer et al., 2022; Dass et al., 2020; Haak, 2025; Invernizzi et al., 2025; Lombardi et al., 2025), with binary DisProt annotations still the main data source used (Mehdiabadi et al., 2026). On the Disorder-PDB challenge of the community-wide CAID assessment, methods have not improved over the last two years (Conte et al., 2023; Mehdiabadi et al., 2026), suggesting that the limit of what is possible with training on binary annotations has been reached.

To investigate the degree to which the performance and generalizability of a prediction method is biased by the training data, we comparatively analyzed the disorder prediction landscape. Rather than seeking to develop a single superior model, we trained a consistent modern architecture on different datasets of protein disorder and flexibility. By creating an evaluation regime without overlap between any training and test set, we could evaluate models trained on all datasets used against all other datasets to investigate how the choice of the training data for IDRs affected performance across a range of disorder and flexibility definitions and datasets.

## 2. Results

To investigate the effect of different disorder definitions on performance, we trained separate UdonPred models on seven curated datasets and evaluated each across all test sets. Performance varied dramatically with no single dataset yielding consistently superior predictions (Figure 2). This demonstrated that annotation disagreements, not model architecture, fundamentall limit predictive accuracy.

**Fig. 1:**
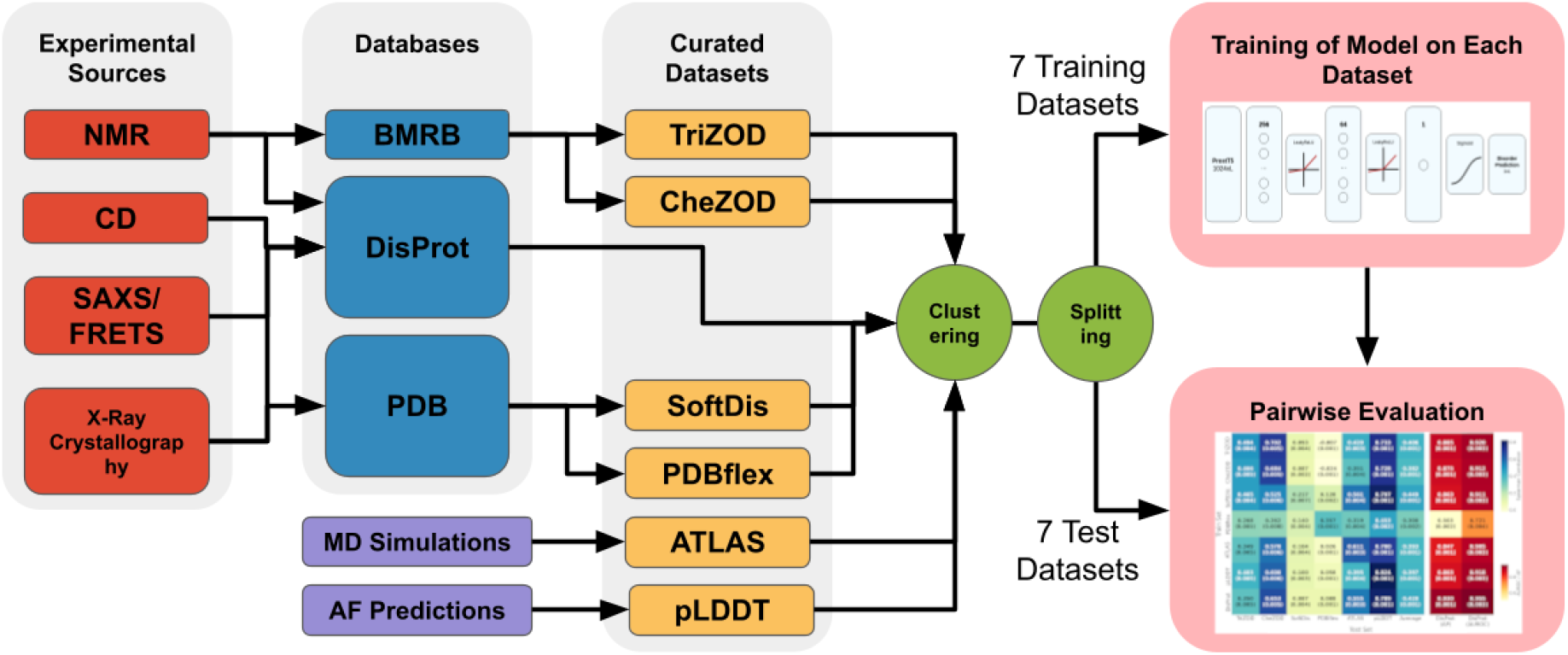
Cross-dataset workflow used to evaluate distinct disorder annotations. Disorder annotations from different experimental sources are deposited in databases and curated into seven distinct datasets representing different definitions of disorder and flexibility. Each dataset is clustered by sequence identity and split into non-redundant training and test sets to prevent data leakage between any train and test datasets. Seven separate UdonPred models - using the same architecture - are trained, one on each training dataset, and evaluated in a pairwise manner against all seven test sets, producing the performance matrix shown in Figure 2.

**Fig. 2:**
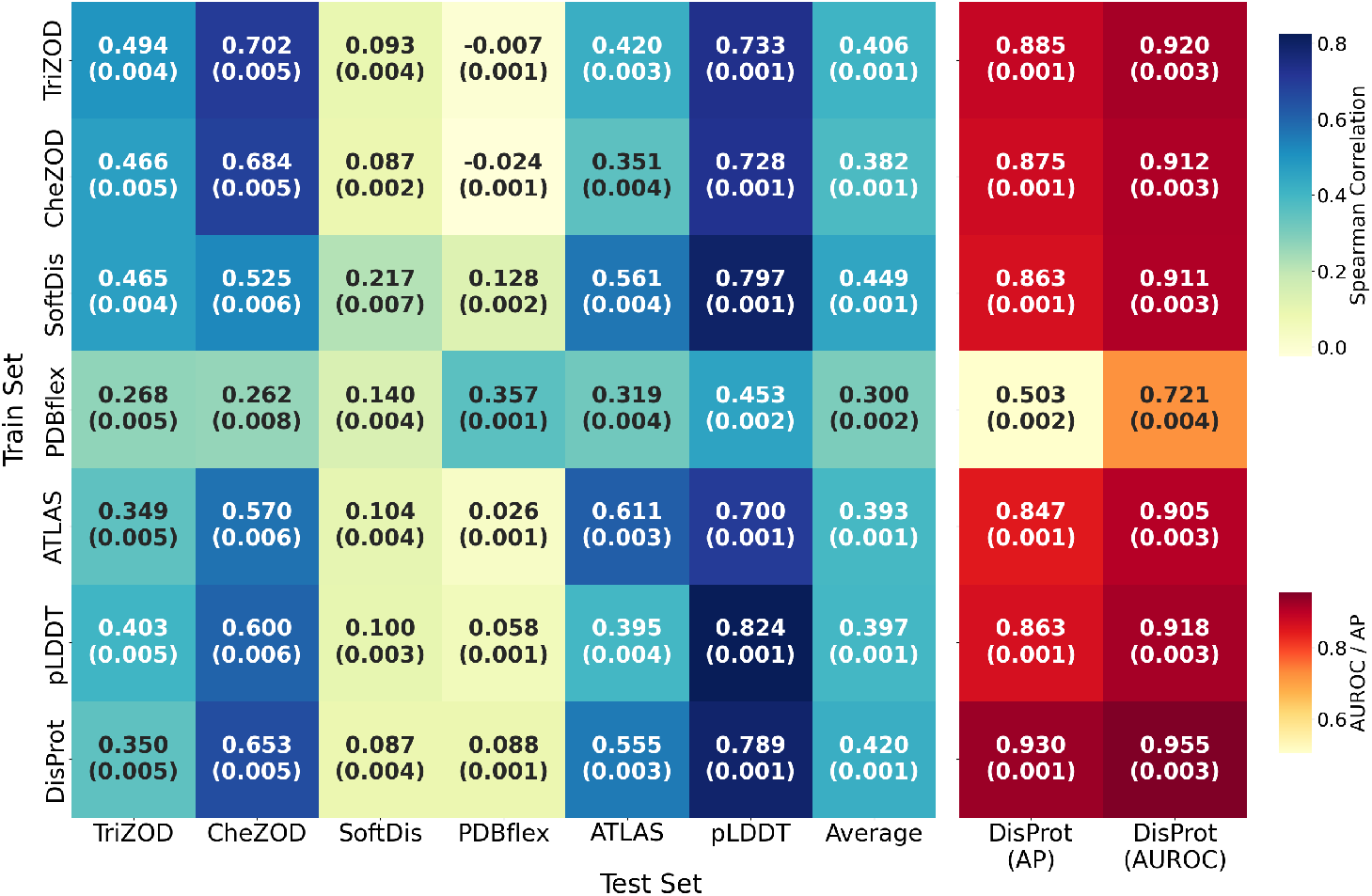
UdonPred performance depends on training dataset choice. Performance matrix of UdonPred models trained on seven different datasets (y-axis) and evaluated on all test sets (x-axis). The left matrix shows Spearman correlations for continuous disorder annotations; the right matrix shows AUROC and Average Precision (AP) for binary DisProt annotations. Diagonal cells represent within-dataset performance. Values in parentheses show 1.96 times the standard error of the mean. Annotation types where higher values indicate lower disorder (CheZOD, pLDDT) are inverted before evaluation.

### 2.1. Cross-dataset performance varies by definition

Flexibility annotations from PDBflex did not work for all datasets. Instead, models trained on PDBflex consistently ranked worst among all methods when evaluated on five of the six other datasets, with SoftDis being the only exception where PDBflextrained models performed second-best. PDBflex annotations are both difficult to predict and inherently noisy. Models trained on other datasets achieved Spearman *r* correlations of 0.128 *±* 0.001 (SoftDis), whereas even training on it directly yielded only 0.357 *±* 0.001.

SoftDis also stood out as hard to learn from other datasets. The best model trained on different data (PDBflex) reached a Spearman correlation of 0.140 *±* 0.004. The limited Spearman correlation of 0.217 *±* 0.006 of the model trained on SoftDis itself suggested SoftDis to be even noisier than PDBflex.

Binary DisProt disorder annotations are seemingly the easiest to learn. Models trained on all datasets except PDBflex reached AP and AUROC scores of *>* 0.84 on the CAID3 test set. The AUROC of the DisProt-trained model was identical (*r* = 0.955 *±* 0.001) to the current best published result on Disorder-PDB, namely PUNCH2 (Meng and Pollastri, 2025; Mehdiabadi et al., 2026). This was unexpected because our DisProt-trained model used a much more simplistic approach than PUNCH2.

Although “only” binary, DisProt annotations appeared informative for other annotations. The model trained on DisProt annotations performed consistently across all test sets except for SoftDis and PDBflex, which all methods predicted less accurately. Model performance varied most for the ATLAS test set (*r* = 0.319 *±* 0.004 trained on PDBflex to 0.511 *±* 0.003 trained on SoftDis).

The distribution of disorder across different datasets appeared less crucial: models trained on datasets with high bias towards ordered residues (TriZOD, ATLAS, PDBflex, SoftDis) did not always perform worse on datasets with less bias (CheZOD, pLDDT, DisProt), and *vice versa*.

Agreement was closest between CheZOD, TriZOD, and pLDDT. The TriZOD model even outperformed the CheZOD model on the CheZOD test set. However, when not considering the pLDDT test set, models trained on these maximally reached a Spearman correlation of 0.702 *±* 0.005 (TriZOD model on CheZOD test set).

No training dataset led to consistently better performance across all test datasets. Every disorder definition and dataset contains nuances that subtly influence the performance of a model trained on it versus other definitions and datasets. The version of `UdonPred` trained on SoftDis exhibited the highest average performance across all test sets, but this is in large part due to it having the only relevant performance on the PDBflex test set, while still performing equally to or slightly lower than other predictors across the other test sets.

### 2.2. Alkylmercury lyase exemplifies per-residue annotation variability

Comparing per-residue annotations of the different datasets for an Alkylmercury lyase that is found in all seven datasets (Fig. 3) serves to illustrate similarities and differences. DisProt annotated two regions as disordered: the first 20 N-terminal residues and a loop from residues 146-161. PDBflex predicted high RMSD values in the N-terminal region marked as disordered by DisProt and low values in the second region compared to even lower values across the rest of the protein, agreeing most with the DisProt annotations. RMSF values in ATLAS varied immensely throughout this protein, with a few local peaks and a pronounced peak at the C-terminus. Inverted pLDDT scores display the same C-terminal peak and agree with the two regions annotated by DisProt. TriZOD, CheZOD and SoftDis all display the pronounced C-terminal peak and peaks in the regions annotated by DisProt (with the exception that TriZOD and CheZOD scores are not available for the N-terminus), but also exhibit more local peaks that are generally not annotated by the other databases, with clearer peaks compared to the background in SoftDis, and noisier peaks in TriZOD and CheZOD that correlate strongly with each other.

**Fig. 3:**
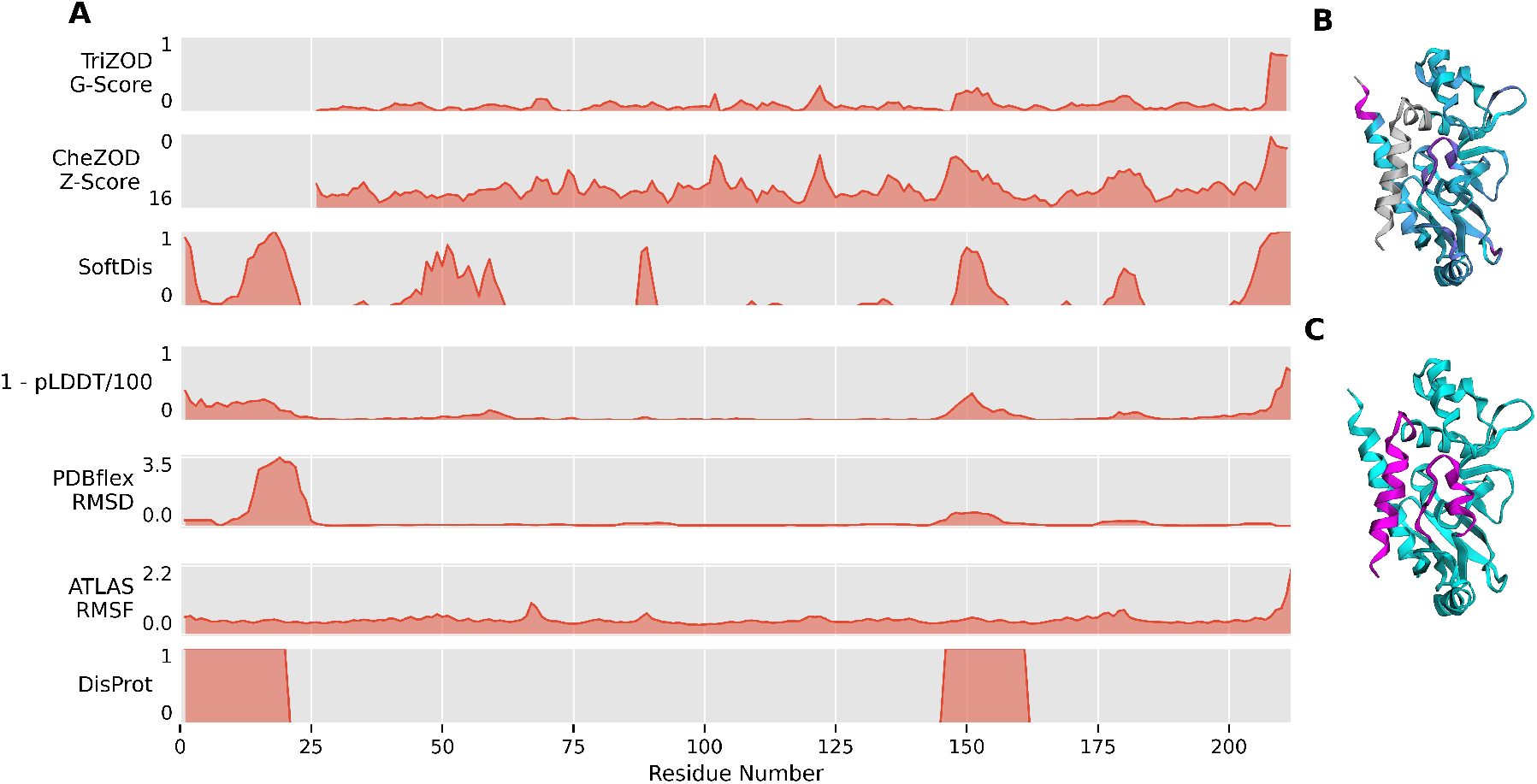
Disorder annotations for the same protein vary substantially across datasets. Case study of Alkylmercury lyase MERB ECOLX (UniProt: P77072). **A** Per-residue disorder annotations from all seven datasets for the 212-residue sequence. **B** AlphaFold2-predicted structure with overlaid continuous TriZOD G-scores (BMRB accession: `6047`). Grey means not annotated, color gradient from turquoise to pink indicates higher degree of disorder annotation. **C** Same structure with overlaid DisProt annotations (accession: `DP00575`). The protein was identified across all datasets through 95% sequence identity clustering. CheZOD and TriZOD lack annotations for the first 25 residues and the final residue. PDBflex and SoftDis sequences contain an extended C-terminus with two additional residues and a His6-tag. The SoftDis sequence has a Ser160Cys substitution, though its cluster includes the canonical sequence due to 95% similarity clustering of cluster representatives.

## 3. Methods

### 3.1. Datasets

#### 3.1.1. DisProt

The DisProt database (Nugnes et al., 2025) is the largest curated database of disorder annotations, containing binary annotations (ordered vs. disordered) for 3.2k proteins. It is widely used for training and evaluating disorder predictors (Wang et al., 2024; Kabir and Hoque, 2024; Hanson et al., 2019; Lombardi et al., 2025) and serves as the reference for CAID experiments (Conte et al., 2023; Mehdiabadi et al., 2026). We used the CAID3 `Disorder-PDB` set, which constrains negative residues to resolved PDB coordinates, excluding “uncertain” residues that lack both structural and disorder annotations, for evaluation. For training, we used the last DisProt release at the time.

#### 3.1.2. CheZOD

CheZOD (Nielsen and Mulder, 2020) (*Chemical Shift Z-score for Order/Disorder*) assigns continuous Z-scores to residues based on NMR chemical shift deviations from random coil predictions calculated from BMRB data (Hoch et al., 2023). Z-scores distinguish ordered from disordered regions using a threshold of 8, providing a more nuanced training signal than binary annotations. CheZOD has been used to train disorder predictors, including ODiNPred (Dass et al., 2020) and SETH (Ilzhöfer et al., 2022). We adhered to the previously defined training set of 1325 proteins and test set of 117 proteins (Dass et al., 2020).

#### 3.1.3. TriZOD

TriZOD (Haak, 2025) extends CheZOD with G-scores, per-residue disorder scores normalized to [0,1] that are independent of chemical shift availability. Unlike CheZOD’s chi-squared deviations, G-scores use the geometric mean of scaled normal distribution density values of backbone NMR chemical shift deviations from random coil predictions (Hoch et al., 2023). TriZOD improves data quality by filtering BMRB datasets based on experimental conditions, excluding non-physiological conditions. The curated dataset contains over 15,000 peptides, representing a 10-fold increase over CheZOD. G-scores were used to train ADOPT2 (Redl et al.; Invernizzi et al., 2025). We used the `tolerant` subset for training and the predefined `Trizod348` set for testing (Haak, 2025).

#### 3.1.4. pLDDT

AlphaFold’s predicted local distance difference test (pLDDT) (Jumper et al., 2021) is a per-residue confidence score (0-100), which inversely correlates with disorder (Piovesan et al., 2022c; Conte et al., 2023), where low scores (pLDDT *<* 50) strongly correlate with intrinsic disorder. We obtained scores for the proteins in DisProt from AlphaFoldDB Varadi et al. (2024) and transformed them to 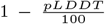 to generate disorder probabilities. We note that AlphaFold 3 has recently been shown to confidently hallucinate structure in disordered regions (Gopalan and Narayanan, 2025).

#### 3.1.5 SoftDis

SoftDis (Seoane and Carbone, 2021; Lombardi et al., 2025) introduces “soft disorder”: residues exhibiting ambiguity between ordered and disordered states through high B-factors or intermittent missing regions across X-ray crystal structures of the same sequence. It is constructed by clustering 500k chains from the Protein Data Bank (PDB; Berman et al., 2000) and contains both classical IDRs (missing regions) and soft disordered regions (SDRs, high B-factors). SoftDis has been used to train the `Lora-DR-Suite` of disorder predictors (Lombardi et al., 2025).

#### 3.1.6. ATLAS

ATLAS defines flexibility using dynamic profiles derived from all-atom Molecular Dynamics (MD) simulations. It characterizes flexibility using metrics like Root Mean Square Fluctuation (RMSF) and entropy-based indices (*N*_*eq*_). It specifically highlights “Dual Personality Fragments” (DPFs), regions that can exist in both ordered and disordered states. The ATLAS dataset consists of standardized MD simulations for representative protein structures from the PDB (Berman et al., 2000) and is curated to ensure exhaustive sampling of conformational space, including specific subsets for chameleon sequences and DPFs. It has been used to train protein ensemble generative models like ESMFlow or FliPS, which also includes a flexibility prediction method, BackFlip (Jing et al., 2024; Viliuga et al., 2025). Since ATLAS is not affected by the same experimental bias as disorder databases and contains information about fine-grained structural dynamics of proteins, it potentially enables a model trained on it to learn broader patterns of structural heterogeneity useful for predicting disorder. We used RMSF scores in training and evaluation.

#### 3.1.7. PDBflex

Similarly, PDBflex (Hrabe et al., 2016) defines flexibility based on structural variations between PDB depositions (Berman et al., 2000) of the same protein chain (*>*95% sequence identity), capturing differences caused by functional states (e.g., apo vs. holo), crystal packing, or intrinsic flexibility. The database clusters about 239k protein chains from the PDB with identical sequences and superimposes these structures to calculate Root Mean Square Deviation (RMSD) and local fluctuation profiles, identifying regions that are unstable or variable across different experimental snapshots. PDBflex has been used to train the protein flexibility predictor MEDUSA (Vander Meersche et al., 2021).

### 3.2. Data processing

We partitioned data into training, validation, and test splits, enforcing *<*30% sequence identity at 80% coverage between splits to prevent data leakage across all datasets.

For the CheZOD, TriZOD, and DisProt datasets, we utilized established benchmark test sets: CheZOD117 (Dass et al., 2020), TriZOD348 (Haak, 2025), and CAID3 Disorder-PDB (Mehdiabadi et al., 2026). For all remaining datasets, test sequences were selected randomly, provided they adhered to the aforementioned sequence identity and coverage constraints against the training and validation splits.

**Table 1.** Number of protein per dataset before and after clustering, as well as split sizes per dataset. TriZOD, CheZOD and DisProt split sizes do not add to the total dataset size. This is because the test split was replaced by a standard test set, with other proteins similar to test proteins being entirely removed.

| Dataset | Total | Clustered | Train | Validation | Test |
| --- | --- | --- | --- | --- | --- |
| TriZOD | 7196 | 7196 | 1027 | 616 | 348 |
| CheZOD | 1291 | 1291 | 235 | 117 | 117 |
| SoftDis | 20267 | 14839 | 8517 | 1278 | 5044 |
| PDBflex | 16017 | 11724 | 7067 | 1164 | 3493 |
| ATLAS | 1390 | 1390 | 802 | 138 | 450 |
| pLDDT | 2566 | 2566 | 1685 | 255 | 626 |
| DisProt | 3150 | 3150 | 1864 | 277 | 319 |

Finally, to optimize computational efficiency during training, the two largest datasets (SoftDis and PDBFlex) were clustered at a 50% sequence identity threshold using MMSeqs2 (Steinegger and Söding, 2017) and only the cluster representatives retained.

### 3.3. Model training

All models use the same UdonPred architecture, which was also submitted to CAID3 (Mehdiabadi et al., 2026) with 1024-dimensional ProstT5 embeddings (Heinzinger et al., 2024) as input. No loss was backpropagated through the language model during training. Disorder prediction is handled by a two-layer feedforward neural network head with two hidden layers of dimensions 256 and 64, respectively. Each layer incorporates a LeakyReLU activation function (Xu et al., 2015) and a dropout rate of 0.4. For datasets with target values constrained to 0-1 (TriZOD, DisProt, and SoftDis), a sigmoid activation function is applied to the final output layer.

Model training was conducted independently on the training split of each dataset with a batch size of 64. The optimization used the AdamW optimizer (Loshchilov and Hutter, 2019) with standard hyperparameters and continued until the loss function reached its minimum on the respective validation set. Specifically, binary cross-entropy (BCE) loss was used for the DisProt dataset, while mean squared error (MSE) was utilized for all other datasets.

### 3.4. Evaluation

The various datasets used here differ in both range and physical interpretation. For instance, TriZOD values are constrained to a [0, 1] interval, representing the deviation of observed chemical shifts from random coil values. In contrast, the ATLAS dataset provides Root Mean Square Fluctuations (RMSF) in Angstroms (Å), which are inherently unbounded. Therefore, direct numerical comparisons between raw predictions and ground-truth values across different datasets are not possible. To address this, we assess monotonic relationships between continuous scores through Spearman rank correlation, and relationships between continuous scores and binary DisProt annotations using area under the receiver-operating curve (AUROC) and average precision (AP), which also facilitate comparison to the CAID3 (Mehdiabadi et al., 2026) benchmark.

## 4. Discussion

### 4.1. Backbone flexibility data enables robust disorder prediction while structural polymorphism does not

Structural polymorphism as defined in PDBflex through the existence of multiple stable, folded conformations in PDB chains is extremely hard to predict for disorder models. Even the model trained on the ATLAS flexibility dataset does not achieve any meaningful performance on the PDBflex test set. The same goes when training models on PDBflex data: their results are moderate, but consistently worse than models trained on other datasets. PDBflex exhibits the strongest case of experimental bias of all datasets used in this work. It is not the only dataset that is exclusively based on X-ray crystallography data (so is SoftDis) or one single experimental data source (so are CheZOD and TriZOD), but crucially, it is biased towards states that can be crystallized and does not contain fragments of more than 20 unidentified residues, which would be annotated as disordered by most intrinsic disorder databases. Having never seen such regions, the UdonPred model trained on PDBflex does not generalize well to them. Using exclusively flexibility data for training is viable, however: training on local flexibility of the protein backbone based on MD simulations from the ATLAS dataset, leads to robust performance on the disorder datasets.

SoftDis, which can be seen as a superset of intrinsic disorder datasets such as DisProt, TriZOD, and CheZOD, exhibits an interesting property: prediction methods trained on other datasets performed much worse on it than the method trained on SoftDis did on those sets. In other words, the loss from training and testing on different data was highest for testing on SoftDis. Possibly, this occurs because SoftDis is a phenomenological superset of intrinsic disorder datasets, as it adds soft disordered regions that are characterized by a high B-factor in X-ray crystallography. This allows the model trained on it to predict disorder and flexibility well, while models trained on more coarse-grained data may fail more because they never learn the fine-grained distinction. However, training on SoftDis also resulted in relatively weak generalization, i.e., relatively low test performance. This might suggest that given the complexity introduced by non-binary classifications in SoftDis, machine learning needs more data for learning the fine-grained distinction.

### 4.2. Intrinsic disorder datasets show agreements despite different annotation sources

Models trained and evaluated on the intrinsic disorder datasets TriZOD, CheZOD, DisProt, and pLDDT transferred well between each other. Training on different annotation types did not lead to consistent differences in correlation on other annotation types, and training on binary disorder annotation types alone did not reduce performance. Additionally, neither size nor distribution of training data consistently impacted performance across different annotations.

Binary DisProt annotations, as well as pLDDT scores of AlphaFold2 structure predictions, however, were predicted with high accuracy from all training sets. Training UdonPred to predict DisProt annotations from sequence alone sufficed to reach the current top performance as identified by the recent CAID3 benchmark `CAID3-PDB`. Similar performance through a much simpler method with substantially fewer over-fittable parameters seems a value on its own. More importantly, it attests to the power of our simple novel solution. While the CAID3 benchmark also contains subsets for disordered binding and linker regions, its disorder challenges might be too easy to challenge modern prediction methods. Therefore future benchmarks would likely benefit from including continuous disorder scores in some form.

In conclusion, still no single type of annotation or database for protein disorder and flexibility presents a better training or evaluation test set than others. At the same time, the rift between different datasets and conflicting definitions of disorder, based on the results presented in this work, seems to be much less limiting than it was just a few years ago. Future methods might be trained in a multimodal capacity to model an expanded representation of disorder dynamics, and future benchmarks should be chosen carefully to include hard samples and counter bias from experimental or data curation techniques in method training.

An obvious limitation of this work is the same as its core strength: we used the same simple set of hyperparameters irrespective of the training data. Thus, we reduced the risk of over-fitting at the expense of possibly improved performance. Hyperparameter tuning, or fine-tuning of a pLM, which has been suggested to lead to good results overall (Schmirler et al., 2024) and specifically for protein disorder (Lombardi et al., 2025), would likely yield improved performance. Combining multiple datasets for training, or more careful sampling during training to account for dataset mismatches and imbalances, are further avenues we did not explore.

## 5. Author Statements

### 5.2. Author contributions statement

T.S. conceptualized and supervised the project. M.H. and T.S. curated TriZOD, CheZOD, and DisProt data. J.S. and D.W. implemented the software and evaluated results on TriZOD, CheZOD, and DisProt under the supervision of T.S., with B.R. and M.H. providing additional feedback. D.W. curated the SoftDis, pLDDT, PDBflex, and ATLAS datasets, expanded the software, and performed further model training and evaluation on them. J.S. drafted and wrote the initial manuscript. J.S., D.W., T.S., and B.R. co-supervised the project and refined the presentation. All authors read and approved the final manuscript.

### 5.3. Use of large language models

Large language models (Gemini, NotebookLM) were used as aids in writing code, correcting written text, and discovering information in accordance with the ISCB Acceptable Use Policy. The authors take full responsibility for all text, references, and code.

## 5.4. Acknowledgments

We thank Nikita Kugut (TUM) for his help with administrative matters, as well as Tobias Olenyi and Lothar Richter (both TUM) for supporting use of the Leibniz Rechenzentrum (LRZ) computing cluster. Our special thanks go out to *Freunde der TUM e*.*V*. for providing funding for D.W. and M.H. to attend the ML4NGP conference in Thessaloniki. We further thank Michael Heinzinger (TUM, Helmholtz Munich) for providing access to the Jülich computing cluster, as well as his indispensable feedback. Our thanks also extend to our colleagues at the TUM and Helmholtz Munich for insightful feedback and discussions. Finally, we thank those who deposit experimental data in public databases, maintain these databases, and develop methods to enrich experimental data.

The work of T.S. and B.R. is supported by the Bavarian Ministry for Education through funding to the TUM. This work was supported by Helmholtz AI computing resources (HAICORE) of the Helmholtz Association’s Initiative and Networking Fund through Helmholtz AI.

## References

M. C. Aspromonte, M. V. Nugnes, F. Quaglia, A. Bouharoua, DisProt Consortium, S. C. E. Tosatto, and D. Piovesan. DisProt in 2024: Improving function annotation of intrinsically disordered proteins. Nucleic Acids Research, 52(D1):D434– D441, Jan. 2024. ISSN 1362-4962. doi: 10.1093/nar/gkad928.

M. M. Babu. The contribution of intrinsically disordered regions to protein function, cellular complexity, and human disease. Biochemical Society Transactions, 44(5):1185–1200, Oct. 2016. ISSN 0300-5127. doi: 10.1042/BST20160172.

M. M. Babu, R. van der Lee, N. S. de Groot, and J. Gsponer. Intrinsically disordered proteins: Regulation and disease. Current Opinion in Structural Biology, 21(3):432–440, June 2011. ISSN 1879-033X. doi: 10.1016/j.sbi.2011.03.011.

H. M. Berman, J. Westbrook, Z. Feng, G. Gilliland, T. N. Bhat, H. Weissig, I. N. Shindyalov, and P. E. Bourne. The Protein Data Bank. Nucleic Acids Research, 28(1):235–242, Jan. 2000. ISSN 0305-1048. doi: 10.1093/nar/28.1.235.

S. E. Bondos, A. K. Dunker, and V. N. Uversky. Intrinsically disordered proteins play diverse roles in cell signaling. Cell Communication and Signaling, 20(1):20, Feb. 2022. ISSN 1478-811X. doi: 10.1186/s12964-022-00821-7.

Y. Cheng, T. LeGall, C. J. Oldfield, A. K. Dunker, and V. N. Uversky. Abundance of Intrinsic Disorder in Protein Associated with Cardiovascular Disease. Biochemistry, 45(35):10448– 10460, Sept. 2006. ISSN 0006-2960. doi: 10.1021/bi060981d.

A. D. Conte, M. Mehdiabadi, A. Bouhraoua, A. Miguel Monzon, S. C. E. Tosatto, and D. Piovesan. Critical assessment of protein intrinsic disorder prediction (CAID) - Results of round 2. Proteins: Structure, Function, and Bioinformatics, 91(12): 1925–1934, 2023. ISSN 1097-0134. doi: 10.1002/prot.26582.

R. Dass, F. A. A. Mulder, and J. T. Nielsen. ODiNPred: Comprehensive prediction of protein order and disorder. Scientific Reports, 10(1):14780, Sept. 2020. ISSN 2045-2322. doi: 10.1038/s41598-020-71716-1.

S. DeForte and V. N. Uversky. Resolving the ambiguity: Making sense of intrinsic disorder when PDB structures disagree. Protein Science, 25(3):676–688, 2016. ISSN 1469-896X. doi: 10.1002/pro.2864.

A. K. Dunker, J. D. Lawson, C. J. Brown, R. M. Williams, P. Romero, J. S. Oh, C. J. Oldfield, A. M. Campen, C. M. Ratliff, K. W. Hipps, J. Ausio, M. S. Nissen, R. Reeves, C. Kang, C. R. Kissinger, R. W. Bailey, M. D. Griswold, W. Chiu, E. C. Garner, and Z. Obradovic. Intrinsically disordered protein. Journal of Molecular Graphics and Modelling, 19(1):26–59, Feb. 2001. ISSN 1093-3263. doi: 10.1016/S1093-3263(00)00138-8.

A. K. Dunker, I. Silman, V. N. Uversky, and J. L. Sussman. Function and structure of inherently disordered proteins. Current Opinion in Structural Biology, 18(6):756–764, Dec. 2008. ISSN 1879-033X. doi: 10.1016/j.sbi.2008.10.002.

A. Elnaggar, M. Heinzinger, C. Dallago, G. Rehawi, Y. Wang, L. Jones, T. Gibbs, T. Feher, C. Angerer, M. Steinegger, D. Bhowmik, and B. Rost. ProtTrans: Toward Understanding the Language of Life Through Self-Supervised Learning. IEEE transactions on pattern analysis and machine intelligence, 44 (10):7112–7127, Oct. 2022. ISSN 1939-3539. doi: 10.1109/TPAMI.2021.3095381.

C. A. Galea, A. High, J. C. Obenauer, A. Mishra, C.-G. Park, M. Punta, A. Schlessinger, J. Ma, B. Rost, C. A. Slaughter, and R. W. Kriwacki. Large-scale Analysis of Thermo-stable, Mammalian Proteins Provides Insights into the Intrinsically Disordered Proteome. Journal of proteome research, 8(1): 211–226, Jan. 2009. ISSN 1535-3893. doi: 10.1021/pr800308v.

S. Gopalan and S. Narayanan. Hallucinations in AlphaFold3 for Intrinsically Disordered Proteins with disorder in Biological Process Residues, Nov. 2025.

T.W. Gräwert and D. I. Svergun. Structural Modeling Using Solution Small-Angle X-ray Scattering (SAXS). Journal of Molecular Biology, 432(9):3078–3092, Apr. 2020. ISSN 0022-2836. doi: 10.1016/j.jmb.2020.01.030.

M. Haak. MarkusHaak/trizod, Dec. 2025.

J. Hanson, K. K. Paliwal, T. Litfin, and Y. Zhou. SPOT-Disorder2: Improved Protein Intrinsic Disorder Prediction by Ensembled Deep Learning. Genomics, Proteomics & Bioinformatics, 17(6):645–656, Dec. 2019. ISSN 1672-0229. doi: 10.1016/j.gpb.2019.01.004.

M. Heinzinger, K. Weissenow, J. G. Sanchez, A. Henkel, M. Mirdita, M. Steinegger, and B. Rost. Bilingual Language Model for Protein Sequence and Structure, Mar. 2024.

J. C. Hoch, K. Baskaran, H. Burr, J. Chin, H. R. Eghbalnia, T. Fujiwara, M. R. Gryk, T. Iwata, C. Kojima, G. Kurisu, D. Maziuk, Y. Miyanoiri, J. R. Wedell, C. Wilburn, H. Yao, and M. Yokochi. Biological Magnetic Resonance Data Bank. Nucleic Acids Research, 51(D1):D368–D376, Jan. 2023. ISSN 0305-1048. doi: 10.1093/nar/gkac1050.

T. Hrabe, Z. Li, M. Sedova, P. Rotkiewicz, L. Jaroszewski, and A. Godzik. PDBFlex: Exploring flexibility in protein structures. Nucleic Acids Research, 44(Database issue):D423–D428, Jan. 2016. ISSN 0305-1048. doi: 10.1093/nar/gkv1316.

C. C. Hsu, M. J. Buehler, and A. Tarakanova. The Order-Disorder Continuum: Linking Predictions of Protein Structure and Disorder through Molecular Simulation. Scientific Reports, 10(1):2068, Feb. 2020. ISSN 2045-2322. doi: 10.1038/s41598-020-58868-w.

L. M. Iakoucheva, C. J. Brown, J. D. Lawson, Z. Obradović, and A. K. Dunker. Intrinsic disorder in cell-signaling and cancer-associated proteins. Journal of Molecular Biology, 323(3):573– 584, Oct. 2002. ISSN 0022-2836. doi: 10.1016/s0022-2836(02)00969-5.

D. Ilzhöfer, M. Heinzinger, and B. Rost. SETH predicts nuances of residue disorder from protein embeddings. Frontiers in Bioinformatics, 2:1019597, 2022. ISSN 2673-7647. doi: 10.3389/fbinf.2022.1019597.

M. Invernizzi, S. Bottaro, J. O. Streit, B. Trentini, N. A. E. Venanzi, D. Reidenbach, Y. Lee, C. Dallago, H. Sirelkhatim, B. Jing, F. Airoldi, K. Lindorff-Larsen, C. Fisicaro, and K. Tamiola. Advancing Protein Ensemble Predictions Across the Order–Disorder Continuum, Nov. 2025. ISSN 2692-8205.

B. Jing, B. Berger, and T. Jaakkola. AlphaFold Meets Flow Matching for Generating Protein Ensembles, Sept. 2024.

J. Jumper, R. Evans, A. Pritzel, T. Green, M. Figurnov, O. Ronneberger, K. Tunyasuvunakool, R. Bates, A. Žídek, A. Potapenko, A. Bridgland, C. Meyer, S. A. A. Kohl, A. J. Ballard, A. Cowie, B. Romera-Paredes, S. Nikolov, R. Jain, J. Adler, T. Back, S. Petersen, D. Reiman, E. Clancy, M. Zielinski, M. Steinegger, M. Pacholska, T. Berghammer, S. Bodenstein, D. Silver, O. Vinyals, A. W. Senior, K. Kavukcuoglu, P. Kohli, and D. Hassabis. Highly accurate protein structure prediction with AlphaFold. Nature, 596(7873):583–589, Aug. 2021. ISSN 1476-4687. doi: 10.1038/s41586-021-03819-2.

M. W. U. Kabir and M. T. Hoque. DisPredict3.0: Prediction of intrinsically disordered regions/proteins using protein language model. Applied Mathematics and Computation, 472:128630, July 2024. ISSN 0096-3003. doi: 10.1016/j.amc.2024.128630.

K. Kotowski, I. Roterman, and K. Stapor. DisorderUnetLM: Validating ProteinUnet for efficient protein intrinsic disorder prediction. Computers in Biology and Medicine, 185:109586, Feb. 2025. ISSN 0010-4825. doi: 10.1016/j.compbiomed.2024.109586.

T. Lazar, A. Connor, C. F. DeLisle, V. Burger, and P. Tompa. Targeting protein disorder: The next hurdle in drug discovery. Nature Reviews Drug Discovery, pages 1–21, June 2025. ISSN 1474-1784. doi: 10.1038/s41573-025-01220-6.

Z. Lin, H. Akin, R. Rao, B. Hie, Z. Zhu, W. Lu, N. Smetanin, R. Verkuil, O. Kabeli, Y. Shmueli, A. dos Santos Costa, M. Fazel-Zarandi, T. Sercu, S. Candido, and A. Rives. Evolutionary-scale prediction of atomic-level protein structure with a language model. Science, 379(6637):1123–1130, Mar. 2023. doi: 10.1126/science.ade2574.

S. Liu, S. Chen, T. Bai, and B. Liu. FusionEncoder: Identification of intrinsically disordered regions based on multi-feature fusion. Bioinformatics, 41(7):btaf362, July 2025. ISSN 1367-4811. doi: 10.1093/bioinformatics/btaf362.

G. Lombardi, B. Seoane, and A. Carbone. LoRA-DR-suite: Adapted embeddings predict intrinsic and soft disorder from protein sequences. Bioinformatics, 41(Suppl 1):i439–i448, July 2025. ISSN 1367-4803. doi: 10.1093/bioinformatics/btaf185.

Loshchilov and F. Hutter. Decoupled Weight Decay Regularization, Jan. 2019.

M. Mehdiabadi, A. Del Conte, M. V. Nugnes, M. C. Aspromonte, S. C. E. Tosatto, and D. Piovesan. Critical Assessment of Protein Intrinsic Disorder Round 3 - Predicting Disorder in the Era of Protein Language Models. Proteins: Structure, Function, and Bioinformatics, 94(1):414–424, 2026. ISSN 1097-0134. doi: 10.1002/prot.70045.

D. Meng and G. Pollastri. PUNCH2: Explore the strategy for intrinsically disordered protein predictor. PLOS ONE, 20(3): e0319208, Mar. 2025. ISSN 1932-6203. doi: 10.1371/journal.pone.0319208.

M. Necci, D. Piovesan, and S. C. E. Tosatto. Where differences resemble: Sequence-feature analysis in curated databases of intrinsically disordered proteins. Database: The Journal of Biological Databases and Curation, 2018:bay127, Dec. 2018. ISSN 1758-0463. doi: 10.1093/database/bay127.

J. T. Nielsen and F. A. A. Mulder. Quality and bias of protein disorder predictors. Scientific Reports, 9(1):5137, Mar. 2019. ISSN 2045-2322. doi: 10.1038/s41598-019-41644-w.

J. T. Nielsen and F. A. A. Mulder. Quantitative Protein Disorder Assessment Using NMR Chemical Shifts. In B. B. Kragelund and K. Skriver, editors, Intrinsically Disordered Proteins: Methods and Protocols, pages 303–317. Springer US, New York, NY, 2020. ISBN 978-1-0716-0524-0. doi: 10.1007/978-1-0716-0524-0_15.

M. V. Nugnes, K. E. A. Bouhraoua, M. Zoubiri, R. Pancsa, E. Fichó, DisProt Consortium, P. Tompa, D. Piovesan, S. C. E. Tosatto, and M. C. Aspromonte. DisProt in 2026: Enhancing intrinsically disordered proteins accessibility, deposition, and annotation. Nucleic Acids Research, page gkaf1175, Nov. 2025. ISSN 1362-4962. doi: 10.1093/nar/gkaf1175.

R. Pancsa and P. Tompa. Structural Disorder in Eukaryotes. PLoS ONE, 7(4):e34687, Apr. 2012. ISSN 1932-6203. doi: 10.1371/journal.pone.0034687.

Y. Pang and B. Liu. DisoFLAG: Accurate prediction of protein intrinsic disorder and its functions using graph-based interaction protein language model. BMC Biology, 22(1):3, Jan. 2024. ISSN 1741-7007. doi: 10.1186/s12915-023-01803-y.

Z. Peng, J. Yan, X. Fan, M. J. Mizianty, B. Xue, K. Wang, G. Hu, V. N. Uversky, and L. Kurgan. Exceptionally abundant exceptions: Comprehensive characterization of intrinsic disorder in all domains of life. Cellular and Molecular Life Sciences, 72(1):137–151, Jan. 2015. ISSN 1420-9071. doi: 10.1007/s00018-014-1661-9.

D. Piovesan, M. Arbesú, M. Fuxreiter, and M. Pons. Editorial: Fuzzy Interactions: Many Facets of Protein Binding. Frontiers in Molecular Biosciences, 9:947215, 2022a. ISSN 2296-889X. doi: 10.3389/fmolb.2022.947215.

D. Piovesan, A. M. Monzon, F. Quaglia, and S. C. E. Tosatto. Databases for intrinsically disordered proteins. Acta Crystallographica Section D: Structural Biology, 78(2):144–151, Feb. 2022b. ISSN 2059-7983. doi: 10.1107/S2059798321012109.

D. Piovesan, A. M. Monzon, and S. C. E. Tosatto. Intrinsic protein disorder and conditional folding in AlphaFoldDB. Protein Science: A Publication of the Protein Society, 31(11):e4466, Nov. 2022c. ISSN 0961-8368. doi: 10.1002/pro.4466.

D. Piovesan, A. Del Conte, D. Clementel, A. M. Monzon, M. Bevilacqua, M. C. Aspromonte, J. A. Iserte, F. E. Orti, C. Marino-Buslje, and S. C. E. Tosatto. MobiDB: 10 years of intrinsically disordered proteins. Nucleic Acids Research, 51 (D1):D438–D444, Jan. 2023. ISSN 0305-1048. doi: 10.1093/nar/gkac1065.

I. Redl, F. Airoldi, S. Bottaro, A. Chung, O. Dutton, C. Fisicaro, P. Foerch, L. Henderson, F. Hoffmann, M. Invernizzi, B. M. J. Owens, S. Ruschetta, and K. Tamiola. Optimizing protein language models with Sentence Transformers.

H. Ruan, Q. Sun, W. Zhang, Y. Liu, and L. Lai. Targeting intrinsically disordered proteins at the edge of chaos. Drug Discovery Today, 24(1):217–227, Jan. 2019. ISSN 1359-6446. doi: 10.1016/j.drudis.2018.09.017.

R. Schmirler, M. Heinzinger, and B. Rost. Fine-tuning protein language models boosts predictions across diverse tasks. Nature Communications, 15(1):7407, Aug. 2024. ISSN 2041-1723. doi: 10.1038/s41467-024-51844-2.

B. Seoane and A. Carbone. The complexity of protein interactions unravelled from structural disorder. PLoS Computational Biology, 17(1):e1008546, Jan. 2021. ISSN 1553-734X. doi: 10.1371/journal.pcbi.1008546.

P. Sormanni, D. Piovesan, G. T. Heller, M. Bonomi, P. Kukic, C. Camilloni, M. Fuxreiter, Z. Dosztanyi, R. V. Pappu, M. M. Babu, S. Longhi, P. Tompa, A. K. Dunker, V. N. Uversky, S. C. E. Tosatto, and M. Vendruscolo. Simultaneous quantification of protein order and disorder. Nature Chemical Biology, 13(4):339–342, Apr. 2017. ISSN 1552-4469. doi: 10.1038/nchembio.2331.

M. Steinegger and J. Söding. MMseqs2 enables sensitive protein sequence searching for the analysis of massive data sets. Nature Biotechnology, 35(11):1026–1028, Nov. 2017. ISSN 1546-1696. doi: 10.1038/nbt.3988.

V. N. Uversky. Intrinsic disorder in proteins associated with neurodegenerative diseases. Frontiers in Bioscience-Landmark, 14(14):5188–5238, June 2009. ISSN 2768-6701. doi: 10.2741/3594.

V. N. Uversky. The triple power of D3: Protein intrinsic disorder in degenerative diseases. Frontiers in Bioscience (Landmark Edition), 19(2):181–258, Jan. 2014. ISSN 2768-6698. doi: 10.2741/4204.

V. N. Uversky. Recent Developments in the Field of Intrinsically Disordered Proteins: Intrinsic Disorder-Based Emergence in Cellular Biology in Light of the Physiological and Pathological Liquid-Liquid Phase Transitions. Annual Review of Biophysics, 50:135–156, May 2021. ISSN 1936-1238. doi: 10.1146/annurev-biophys-062920-063704.

V. N. Uversky, V. Davé, L. M. Iakoucheva, P. Malaney, S. J. Metallo, R. R. Pathak, and A. C. Joerger. Pathological Unfoldomics of Uncontrolled Chaos: Intrinsically Disordered Proteins and Human Diseases. Chemical reviews, 114(13): 6844–6879, July 2014. ISSN 0009-2665. doi: 10.1021/cr400713r.

Y. Vander Meersche, G. Cretin, A. G. de Brevern, J.-C. Gelly, and T. Galochkina. MEDUSA: Prediction of Protein Flexibility from Sequence. Journal of Molecular Biology, 433(11):166882, May 2021. ISSN 0022-2836. doi: 10.1016/j.jmb.2021.166882.

M. Varadi, D. Bertoni, P. Magana, U. Paramval, I. Pidruchna, M. Radhakrishnan, M. Tsenkov, S. Nair, M. Mirdita, J. Yeo, O. Kovalevskiy, K. Tunyasuvunakool, A. Laydon, A. Žídek, H. Tomlinson, D. Hariharan, J. Abrahamson, T. Green, J. Jumper, E. Birney, M. Steinegger, D. Hassabis, and S. Velankar. AlphaFold Protein Structure Database in 2024: Providing structure coverage for over 214 million protein sequences. Nucleic Acids Research, 52(D1):D368–D375, Jan. 2024. ISSN 0305-1048. doi: 10.1093/nar/gkad1011.

V. Viliuga, L. Seute, N. Wolf, S. Wagner, A. Elofsson, J. Stühmer, and F. Gräter. Flexibility-Conditioned Protein Structure Design with Flow Matching, June 2025.

K. Wang, G. Hu, S. Basu, and L. Kurgan. flDPnn2: Accurate and Fast Predictor of Intrinsic Disorder in Proteins. Journal of Molecular Biology, 436(17):168605, Sept. 2024. ISSN 0022-2836. doi: 10.1016/j.jmb.2024.168605.

P. E. Wright and H. J. Dyson. Intrinsically unstructured proteins: Re-assessing the protein structure-function paradigm. Journal of Molecular Biology, 293(2):321–331, Oct. 1999. ISSN 0022-2836. doi: 10.1006/jmbi.1999.3110.

B. Xu, N. Wang, T. Chen, and M. Li. Empirical Evaluation of Rectified Activations in Convolutional Network, Nov. 2015.

B. Xue, A. K. Dunker, and V. N. Uversky. Orderly order in protein intrinsic disorder distribution: Disorder in 3500 proteomes from viruses and the three domains of life. Journal of Biomolecular Structure and Dynamics, 30(2):137–149, June 2012. ISSN 0739-1102. doi: 10.1080/07391102.2012.675145.

X. Zhang, X. Song, G. Hu, Y. Yang, R. Liu, N. Zhou, S. Basu, D. Qiao, and Q. Hou. Landscape of intrinsically disordered proteins in mental disorder diseases. Computational and Structural Biotechnology Journal, 23:3839–3849, Dec. 2024. ISSN 2001-0370. doi: 10.1016/j.csbj.2024.10.043.

